# Genetic characterization and dissemination of *Staphylococcus aureus* and Staphylococci genus: food and health perspective

**DOI:** 10.1101/2023.07.24.550257

**Authors:** Dalal M. Alkuraythi, Manal M. Alkhulaifi, Abdulwahab Z. Binjomah, Mohammed Alarawi, Hind M Aldakhil, Mohammed I. Mujallad, Saleh Ali Alharbi, Mohammed Alshomrani, Saeed Mastour Alshahrani, Takashi Gojobori, Sluiman M Alajel

## Abstract

Hospitals and communities are significant hubs for antibiotic-resistant staphylococcal infections, but recently there have been spread of the Staphylococci to other settings including farms and food chains. This study aimed to compare the methicillin-resistant *Staphylococcus aureus* (MRSA) isolates from humans and food to understand the genetic characteristics and similarities between the circulating clones across both groups. A total of 250 samples of meat (camel, beef, chicken, fish, and lamb) were collected from different retailers in Riyadh, Saudi Arabia. Culture, and Genomics analysis of the isolates yielded 53 *S*.*aureus* out of which (42/53; 79%) were methicillin-sensitive *S*.*aureus* (MSSA) and (11/53; 21%) were MRSA in addition to to other Staphylococci. 80 clinically confirmed MRSA isolates were obtained from patients in the same city. The most common *S. aureus* clone in patients and retail meat belonged to clonal complex (CC5). In meat, ST6 and ST97 were the most common clones found in (6/11; 54.5%) of MRSA isolates, and ST1153 (9/42; 21.42%), ST672 (7/42; 16.66%) of MSSA isolates. The majority of MRSA isolates in meat carried SCC*mec* type V, which was observed in (6/11; 54.54%) of isolates. While in patients, ST5 and ST6 were the predominant clones found in (37/80; 46%) of MRSA isolates. The most prevalent SCC*mec* types in MRSA isolates from patients were SCC*mec* type IVa, which was observed in (39/80; 48.7%) of isolates. While there was no MRSA found in beef, camel meat had the highest prevalence of MRSA ST6-t2450 contamination. The other two clones, CC97 and CC361, were the second-most prevalent clones in meat and were relatively common among patients. Novel *S. aureus* lineages were sequenced and characterized for the first time ST8109 from meat and ST8110 and ST8111 from patients. A structured One Health approach on *S. aureus* considering its dissemination, genetic characterization, antibiotic resistance, and overall impact on food intended for human consumption and its effect on human health is advised.

## Introduction

*Staphylococcus aureus* is a gram-positive commensal bacterium that is widespread and often inhabits several human and animal body parts, primarily the nostrils and skin [1]. It is generally harmless bacteria, but an opportunistic pathogen and may cause a wide range of diseases, that varies from mild skin infections to life-threatening infections such as bacteremia and sepsis [2]. Early in the 1960’s, *S. aureus* methicillin resistance was observed, following the introduction of semi-synthetic beta-lactams, which are thought to overcome penicillin’s growing resistance, which resulted in further complications of the treatment of *S. aureus* infections [3, 4]. In Saudi Arabia, the prevalence of methicillin resistant *S. aureus* (MRSA) among hospital infections is increasing, and the country’s average MRSA infection rate is as high as 38% [6]. Animals raised for food are also affected by the antimicrobial-resistant bacteria that emerged from the abuse of antibiotics, therefore the food chain sectors are affected as well [7]. Livestock-associated MRSA (LA-MRSA), which is represented mainly by clonal complex 398 (CC398), is the main MRSA reservoir outside of hospitals, colonizing calves, cows, sheep, and poultry [8, 9]. While several hospital-acquired MRSA (HA-MRSA) and community-associated MRSA (CA-MRSA) lineages are known to cause infections in humans, some of them, including CC1, CC5, CC8, CC9, CC30, CC22, and CC97, have become successful LA-MRSA clones [10]. Due to its clinical significance, researchers in Saudi Arabia are focused more on HA-MRSA than other types of MRSA [11]. Even though several studies have been conducted in Saudi Arabia which address the molecular characterization of MRSA [12–15], the volume of data available about MRSA and Staphylococci genetic composition in Saudi Arabia is limited [16]. For that reason, comparative genomics analysis of the genetic make-up of various clones of MRSA isolates obtained from humans and food is needed, also MRSA surveillance at the human-animal-food interfaces must be maintained [17].

## Materials and methods

### Samples collection

Eighty-four non-duplicating clinically identified *S. aureus* routine cultures were sourced from the bacteriology department in Riyadh Regional Laboratory and Blood Bank (RRLBB), various types of analyzed specimens from patients were collected from February to June 2022. 250 retail meat samples (camel, beef, chicken, fish, and lamb) were collected from October 2021 to March 2022 from various meat retailers throughout the city of Riyadh, Saudi Arabia. All meat samples were transported in sealed plastic wrap or the original packaging to the Reference Laboratory of Microbiology at the Saudi Food and Drug Authority (SFDA). Samples were subjected to microbiological analysis within 24 hours after sampling.

### Identification and confirmation of *S. aureus* and MRSA isolates

All obtained clinical isolates were identified as MRSA using VITEK2 (bioMérieux, France) and BD Phoenix (BD Diagnostics, USA). To homogenize the meat samples, a sterile plastic bag with 25 grams of each meat sample and 225 ml of buffered peptone water (BPW) containing 6.5% of NaCl was sealed and homogenized in a stomacher for 2 minutes. Then, the Petrifilm™ Staph Express count plate (3M™, USA) was used after serial dilutions and incubated for 24 hours. Then Petrifilm™ Staph Express Disk was used to distinguish *S. aureus* colonies from other Staphylococci, single *S. aureus* colonies were transferred on mannitol salt agar (MSA) (Neogen, USA). *S. aureus* was identified and confirmed by the classical method, gram stain, catalase test, and coagulase test (PROLEX™, UK). All *S. aureus* isolates were identified by Matrix-Assisted Laser Desorption/Ionization Time-of-Flight mass-spectrometer (MALDI-TOF) (BRUKER, Germany) by 70% formic acid protein extraction method.

### Culture-based methods

MRSA Chromogenic Agar was used for the selective and differential detection of MRSA. Single colonies of *S. aureus* isolates were transferred to Harlequin MRSA Chromogenic agar (NEOGEN, USA). Cefoxitin (Fox), a supplement to this chromogenic medium, prevents the growth of methicillin-sensitive *S. aureus* (MSSA) and allows the growth of MRSA. The *α*-glucosidase produced by *S. aureus* cleaves the chromogenic substrate and gives a blue color to the colonies.

### Molecular-based methods

DNA was extracted from the bacterial culture using QIAGEN DNeasy blood and tissue kit (Qiagen, England, UK) as per the manufacturer’s instructions, DNA purity was checked using QIAxpert spectrophotometer device (Qiagen, England, UK), DNA concentration was determined using Qubit™ Flex Fluorometer device (ThermoFisher Scientific, USA). Detection of the methicillin resistance was determined using the *mecA* gene encoding for the Penicillin’s binding protein 2a (PBP2a), using forward primer 5’ GTAGAAATGACTGAACGTCCGATAA-3’ and reverse 5’ CCAATTCCACATTGTTTCCGTCTAA-3’ amplifying the 310 bp fragment [18]. The reaction was performed in 25uL using DreamTaq™ 2X Green PCR master mix (Thermo Scientific, USA). The reaction was set using 1uL of each (10 pmol) primer, and 2uL of DNA template (20 ng/uL). *S. aureus* ATCC43300 was used as a positive control (ATCC, USA). The amplification was carried out as follows: initial denaturation at 95C° for 4 minutes, followed by 30 cycles of 95C° for 45 seconds, 56C° for 1 minute, 72C° for 1 minute, and a final extension of 72C° for 4 minutes. The final PCR product was analyzed on 1.6% agarose gel stained with Ethidium bromide running 60 minutes at 80 volts in 1X tris borate buffer (TBE) (BIOBasic, CA). The gel visualization was done using Syngene image analysis software (Syngene, USA).

### Whole genome sequencing

Whole genome shotgun (WGS) libraries were constructed using a QIAseqFX DNA library preparation kit, the input concentration was 100 ng of DNA following the manufacturer’s protocol (Qiagen, UK). The library was size selected to 300-350bp insert size. Sequencing was performed using Illumina Novaseq 6000 platform using two SP lanes. Data were quality-checked prior to analysis, and the Phred score cutoff was Q30. The bioinformatics analysis was conducted using the BactopiaV2 pipeline with a focus on *S. aureus* specific workflows [19]. Screening for Panton–Valentine Leukocidin (PVL) was carried out using locally created blast database identification of the two genes lukS and LukF, NCBI accessions (YP 002268030.1, and YP 002268029.1) respectively. For the sequence type (ST) assignment, novel ST sequences were submitted to https://pubmlst.org/saureus/

### Statistical analysis

The frequency of the species, sequence type, and *mecA* by the source of the isolates (either patients or meat) were assessed using SPSS 21.0 statistical software (SPSS Inc., Chicago, IL, USA). The association between species, sequence type, and *mecA* by the source was evaluated using Chi-square Fisher’s exact test with using *α*= 0.05 as the level of significance for this analysis.

## Results

In Riyadh city, 160 positive Staphylococci isolates were isolated from patients and meat Figure(1). Regarding the sequence type, the most common sequence types of *S. aureus* isolates from patients were ST5, ST6, and ST88, whereas the most common sequence types of *S. aureus* isolates from meat were ST672 and ST1153 (*P ≤* 0.001). A total of 76 Staphylococci isolates from meat were identified, including camel (23/76; 30%), fish (15/76; 20%), beef (14/76; 18%), chicken (13/76; 17%), and lamb (11/76; 15%). Fifty-three of the 76 food-isolated Staphylococci were *S. aureus*, of which (42/53; 79%) were MSSA and (11/53; 21%) were MRSA. No significant difference was detected between MRSA prevalence among the different meat types (*P* ≥ 0.05). The remaining Staphylococci (23/76; 30%) belonged to other species table(1). The highest contaminated type of meat with MRSA was camel meat (6/11; 54%), and the lowest contaminated type of meat was lamb and fish (1/11; 9%), While no MRSA in beef was detected. MSSA was detected equally in camel and beef meat (12/42; 28%), and chicken was the least type of meat contaminated with MSSA (4/42; 10%). Fish meat was the most contaminated with other Staphylococci species (7/23; 30%), whereas beef was the least contaminated (2/23; 9%) with other Staphylococci species. In patients, (80/84; 95%) of the obtained clinical isolates were MRSA, the rests were MSSA isolates, and one *S. argentous* isolate.

**Fig 1.**
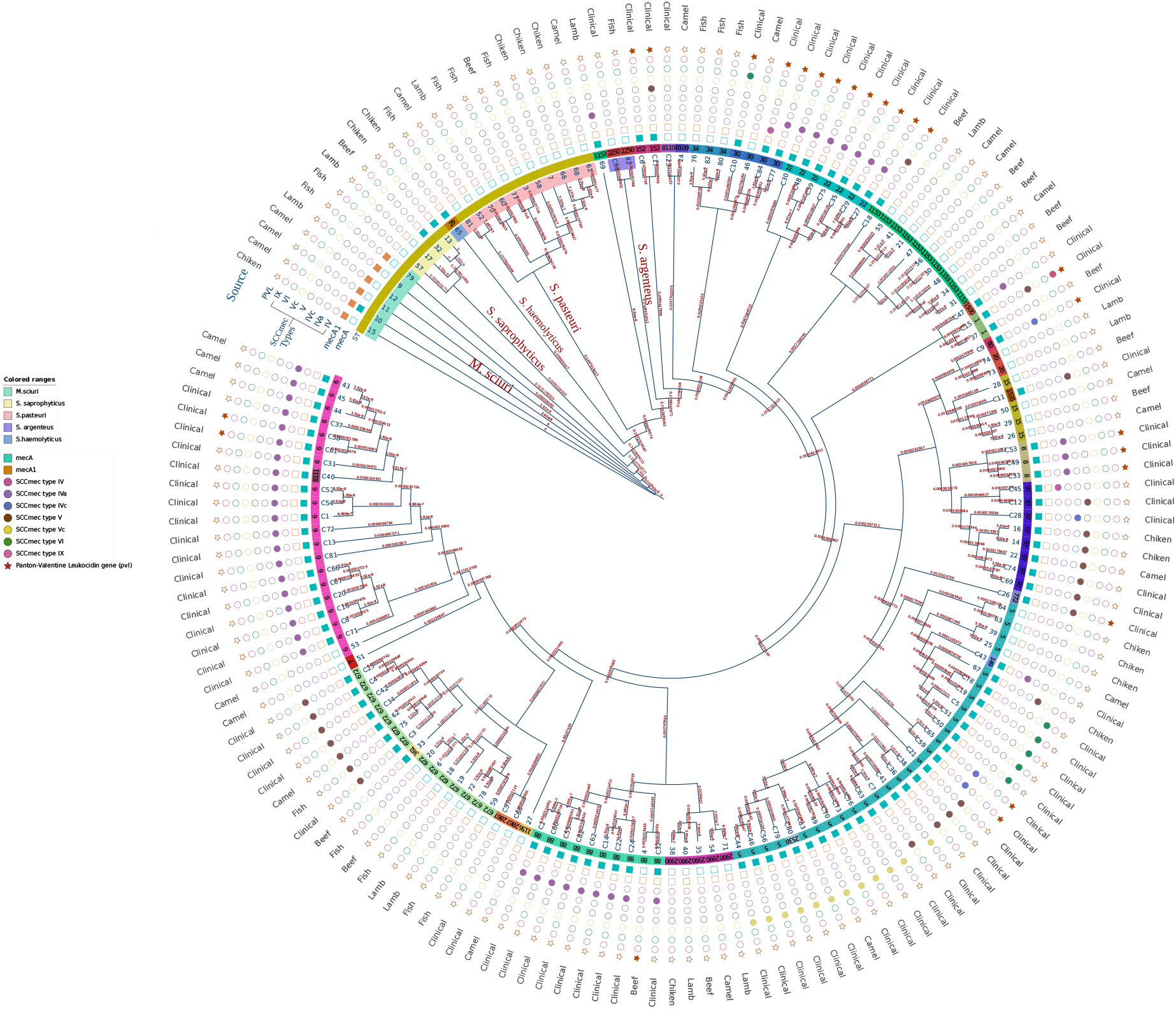
Staphylococci phylogenetic tree: A total of 160 Staphylococci isolates, including *S. aureus* in 15 clonal complexes (CC) were used to create a SNPs based core genome phylogeny. a. Inner colored ranges of the circle indicate other species of Staphylococci. b. Shapes outside the circle indicates the presence of *mecA* and *mecA*1 genes, types of SCC*mec*, and pvl gene presence.

**Table 1.**
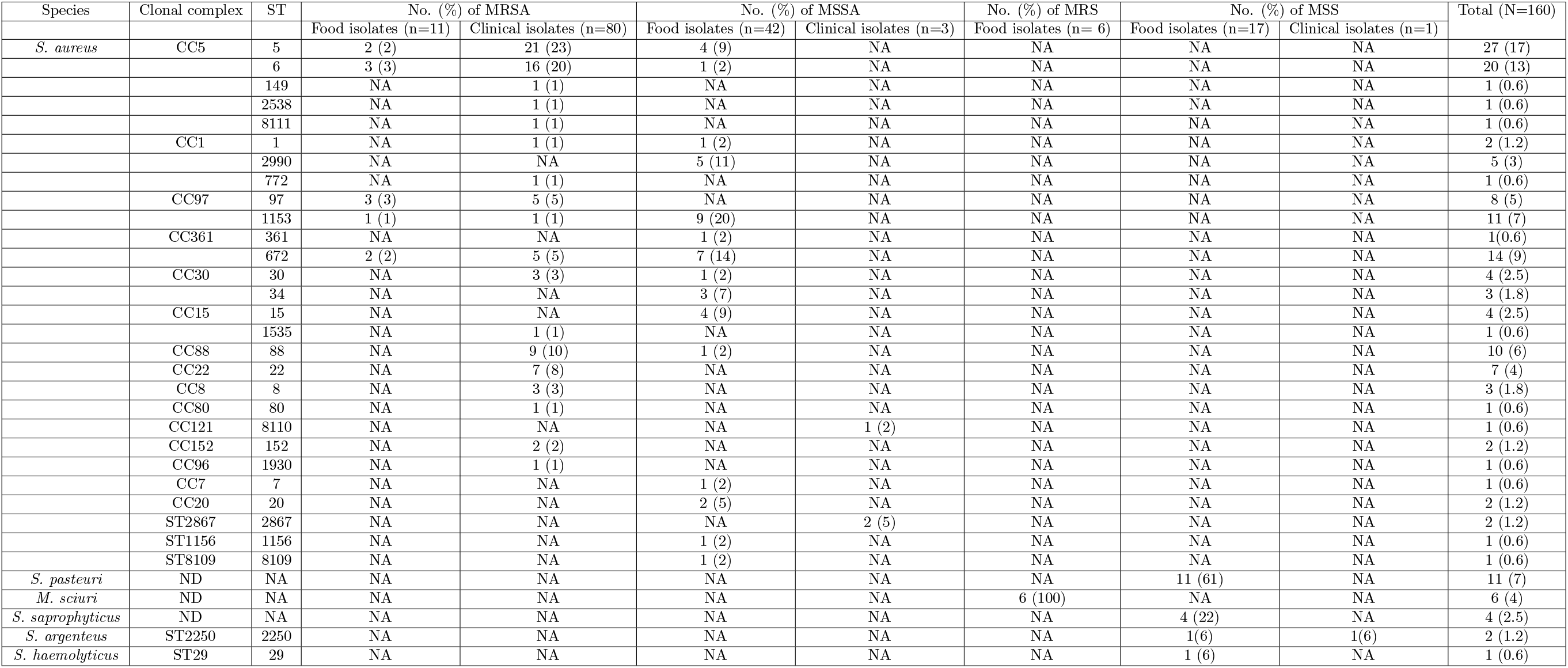
MRSA and MSSA clonal lineages and sequence types (STs) among *S. aureus* isolates from meat and patients, n, the number of the total isolates from the type of sample; N, The total number of *S. aureus* isolates; NA, no isolates positive for this sequence type; ND, the clonal lineages for this strain not detectable; Numbers between parentheses indicate the percentage

### *S. aureus* detection based on MALDI-TOF-MS

Utilizing conventional identification for meat isolates with the MALDI-TOF MS technology, (73/76; 96%) strains belonging to 6 distinct species from the Staphylococci genus were identified. Of which (52/76; 68%) were identified as *S. aureus*, (8/76; 11%) were identified as *S. pasteuri*, (6/76; 8%) were identified as *S. sciuri*, (3/76; 4%) were identified as *S. saprophyticus*, (2/76; 3%) were identified as *S. warneri*, (1/76; 1%) was identified as *S. haemolyticus*, (1/76; 1%) was identified as *S. argentous* and (3/76; 4%) strains were not identified with no peaks detected.

### PCR detection of *mecA* gene

All 76 meat Staphylococci isolates were subjected to *mecA* gene detection by conventional PCR. Among the 53 *S. aureus* isolates from meat, 11 were MRSA, *mecA* gene positive, and 42 were MSSA, *mecA* gene negative.

### Automated MRSA detection

VITEK2 and BD Phoenix™ automated system: VITEK2 was used to identify and confirm 57 *S. aureus* isolates from patients as MRSA. Fifty-six isolates were cefoxitin screen positive and had an oxacillin MIC of 4 *≥* mg/liter. One isolate was cefoxitin screen negative and had an oxacillin MIC of 0.5 mg/liter. BD Phoenix™ was used to identify and confirm 26 isolates from patients as MRSA. All isolates had a cefoxitin MIC of *≥* 8 mg/liter and an oxacillin MIC of *≥* 2 mg/liter.

### Molecular types of Staphylococci isolates from meat

Multilocus sequence typing (MLST) analysis revealed a total of 16 STs of *S. aureus* isolated from meat Figure(2a). Five different STs were assigned to 11 MRSA isolates from meat: ST5, ST6, ST97, ST1153, and ST672 table (1). ST6 and ST97 were the most common clones found (6/11; 54.5%) of MRSA isolates. The remaining clones belonged to ST5 (2/11; 18.18%), ST1153 (1/11; 9%), and ST672 (2/11; 18%). These clones belonged to clonal complexes, CC5 (ST5 and ST6), CC97 (ST97 and ST1153), and CC361 (ST672). In contrast, MSSA isolates from meat were assigned to 15 STs: ST5, ST6, ST1, ST2990, ST1153, ST361, ST672, ST30, ST34, ST15, ST88, ST7, ST20, ST1156, and a novel sequence type ST8109 assigned as a result to this study table(1); Figure(2,b). The predominant clones within the MSSA strains were ST1153 (9/42; 21.42%), ST672 (7/42; 16.66%), and ST2990 (5/42; 11.9%) (Figure 3). The MSSA clones belonged to CCs: CC5 (ST5 and ST6), CC1 (ST1and ST2990), CC97 (ST1153), CC361(ST361, ST672), CC30 (ST30 and ST34), CC15 (ST15), CC88 (ST88), CC7 (ST7), and CC20 (ST20). Methicillin-resistant Staphylococci (MRS) were present among meat isolates, whereas all six isolates belonged to *Mammaliicoccus sciuri* (previously known as *Staphylococcus sciuri*) and harbored the *mecA*1 gene. Methicillin-susceptible Staphylococci (MSS) were also detected in meat isolates, *Staphylococcus pasteuri* (11/23; 47.82%), *Staphylococcus saprophyticus* (4/23; 17.39%), and one isolate from each of *Staphylococcus argenteus* ST2250 and *Staphylococcus haemolyticus* ST29 (2/23; 8.69%).

**Fig 2.**
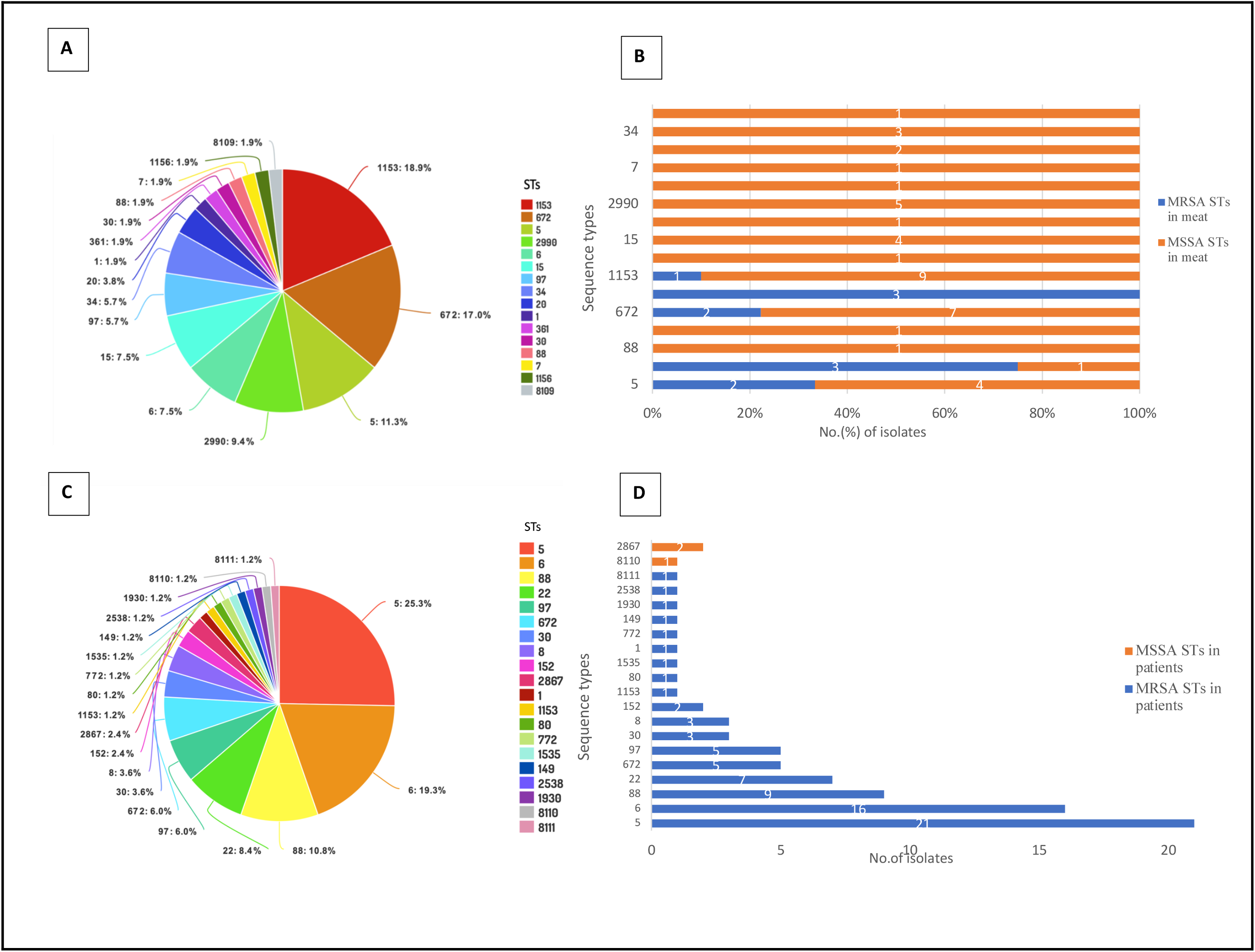
MLST typing results of *S. aureus* in this study. a. The distribution of *S. aureus* STs in meat, b. The distribution of MSSA and MRSA STs in meat, c. The distribution of *S. aureus* STs in patients, d. The distribution of MSSA and MRSA STs in patients.

**Fig 3.**
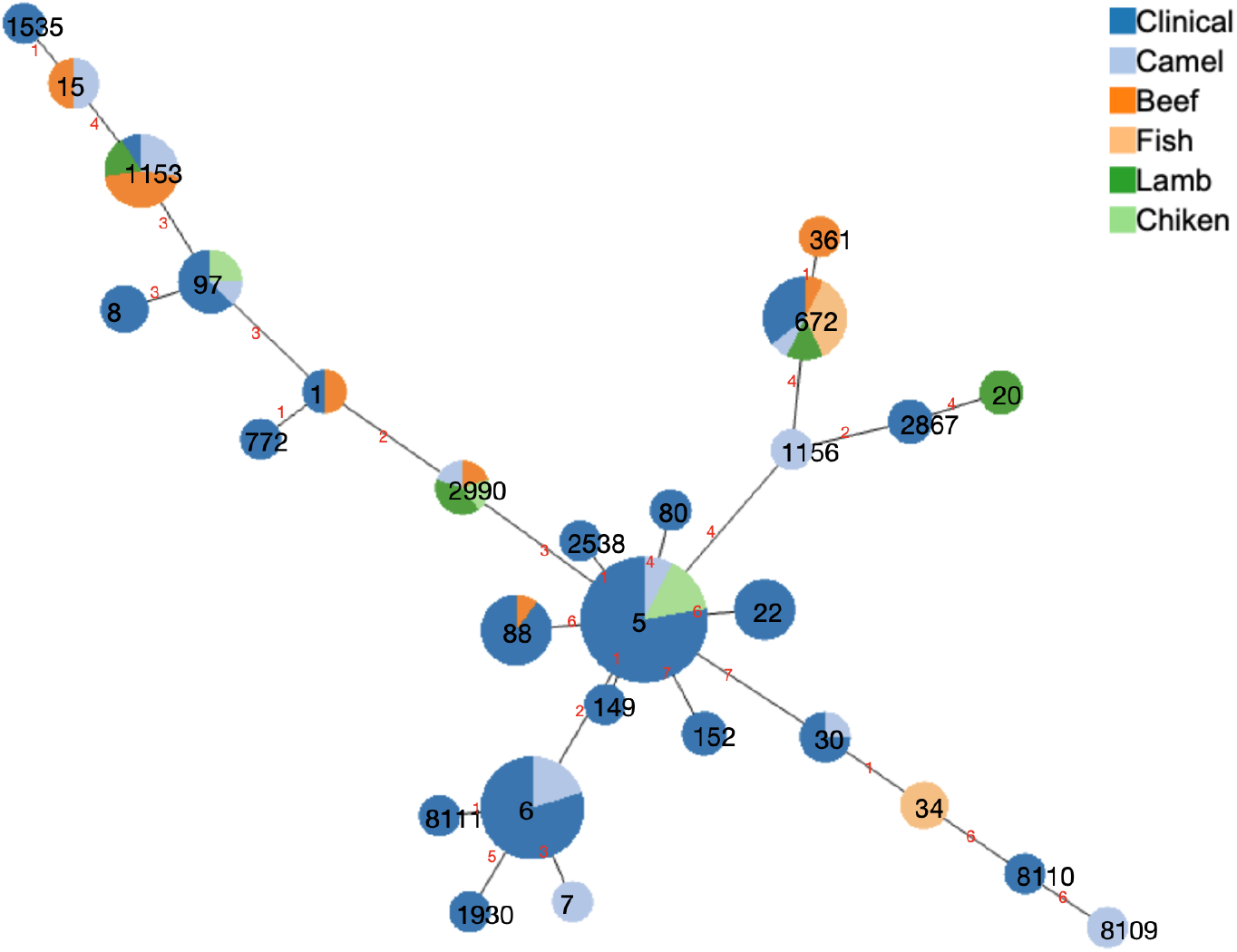
Minimal Spanning Tree (MST) was created with PHYLOViZi [5]: Analysis of *S. aureus* clones based on MLST data. Each circle corresponds to a ST. The size of each circle corresponds to the number of isolates that belong to the ST.

### SCC*mec* types and *spa* types of *S. aureus* from meat

SCC*mec* types and *spa* types of *S. aureus* from meat : The majority of MRSA isolates carried SCC*mec* type V, which was observed in (6/11; 54.54%) of isolates, followed by SCC*mec* type IVa, which was present in (4/11; 36.36%) of the isolates, and one isolate harboring SCC*mec* type Vc figure(1). MRSA isolates had 7 *Staphylococcus aureus* protein A *spa* types, t12375, t903, t311, t2450, t688, t434, and t3841 which were relatively equally distributed across MRSA isolates table(2). Despite this, certain types of meat have more frequent clones, such as MRSA ST6-t2450 in camel meat and ST97-t12375 in chicken. MSSA isolates yielded 17 *spa* types and one unknown *spa* type. The most predominant *spa* types were t003, t903, and t091, which were associated with clones, ST672, ST1153, and ST2990 accounting for 52.38% of all *spa* types in MSSA isolates. Other *spa* types detected in fewer MSSA isolates were t13275, t442, t084, t213, t605, t127, t1339, t3519, t6047, t304, t3478, t164, and t166 table(2).

**Table 2.** Frequency of *spa* types and related STs of *S. aureus* isolates from meat and patients. NA, no isolates positive for this *spa* type; Numbers between parentheses indicate the percentage11/19

| *spa* type | ST | Frequency | *mecA* |  | Isolation source |  |
| --- | --- | --- | --- | --- | --- | --- |
|  |  |  | MSSA | MRSA | Meat | Patients |
| t304 | 6 | 15 (11.4%) | 1 | 14 | 1 | 14 |
| t311 | 5 | 13 (9.9%) | NA | 13 | 1 | 12 |
| t903 | 1153 | 10 (7.6%) | 8 | 2 | 9 | 1 |
| t003 | 672, 361 | 8 (6.1%) | 8 | NA | 8 | NA |
| t3841 | 672 | 7 (5.3%) | NA | 7 | 2 | 5 |
| t091 | 2990, 7 | 6 (4.5%) | 6 | NA | 6 | NA |
| t688 | 5 | 6 (4.5%) | NA | 6 | 1 | 5 |
| t690 | 88 | 5 (3.8%) | NA | 5 | NA | 5 |
| t005 | 22 | 4 (3.0%) | NA | 4 | NA | 4 |
| t2450 | 6 | 3 (2.2%) | NA | 3 | 3 | NA |
| t084 | 15, 1535 | 3 (2.2%) | 2 | 1 | 2 | 1 |
| t166 | 34 | 3 (2.2%) | 3 | NA | 3 | NA |
| t309 | 22 | 3 (2.2%) | NA | 3 | NA | 3 |
| t008 | 8 | 3 (2.2%) | NA | 3 | NA | 3 |
| t267 | 97 | 3 (2.2%) | NA | 3 | NA | 3 |
| t12375 | 97, 8109 | 2 (1.5%) | 1 | 2 | 3 | NA |
| t3478 | 5 | 2 (1.5%) | 2 | NA | 2 | NA |
| t442 | 5 | 2 (1.5%) | 2 | NA | 2 | NA |
| t127 | 1 | 2 (1.5%) | 1 | 1 | 1 | 1 |
| t164 | 20 | 2 (1.5%) | 2 | NA | 2 | NA |
| t105 | 5 | 2 (1.5%) | NA | 2 | NA | 2 |
| t021 | 30 | 2 (1.5%) | NA | 2 | NA | 2 |
| t434 | 97 | 1 (0.7%) | NA | 1 | 1 | NA |
| t213 | 1156 | 1 (0.7%) | 1 | NA | 1 | NA |
| t605 | 15 | 1 (0.7%) | 1 | NA | 1 | NA |
| t3519 | 30 | 1 (0.7%) | 1 | NA | 1 | NA |
| t6047 | 15 | 1 (0.7%) | 1 | NA | 1 | NA |
| t1339 | 88 | 1 (0.7%) | 1 | NA | 1 | NA |
| t355 | 152 | 1 (0.7%) | NA | 1 | NA | 1 |
| t018 | 30 | 1 (0.7%) | NA | 1 | NA | 1 |
| t2297 | 97 | 1 (0.7%) | NA | 1 | NA | 1 |
| t4019 | 152 | 1 (0.7%) | NA | 1 | NA | 1 |
| t159 | 8110 | 1 (0.7%) | 1 | NA | NA | 1 |
| t345 | 772 | 1 (0.7%) | NA | 1 | NA | 1 |
| t521 | 97 | 1 (0.7%) | NA | 1 | NA | 1 |
| t2177 | 88 | 1 (0.7%) | NA | 1 | NA | 1 |
| t9736 | 8111 | 1 (0.7%) | NA | 1 | NA | 1 |
| t045 | 149 | 1 (0.7%) | NA | 1 | NA | 1 |
| t4570 | 1930 | 1 (0.7%) | NA | 1 | NA | 1 |
| t2016 | 2867 | 1 (0.7%) | 1 | NA | NA | 1 |
| t2649 | 88 | 1 (0.7%) | NA | 1 | NA | 1 |
| t1627 | 6 | 1 (0.7%) | NA | 1 | NA | 1 |
| t4494 | 88 | 1 (0.7%) | NA | 1 | NA | 1 |
| t8657 | 6 | 1 (0.7%) | NA | 1 | NA | 1 |
| t3778 | 2538 | 1 (0.7%) | NA | 1 | NA | 1 |
| t12438 | 88 | 1 (0.7%) | NA | 1 | NA | 1 |
| t319 | 5 | 1 (0.7%) | NA | 1 | NA | 1 |

### Molecular types of Staphylococci isolates from patients

According to MLST analysis, 20 STs were assigned *S. aureus* isolates from patients figure(2.c). A total of 80 MRSA isolates from patients were assigned to 18 distinct STs: ST5, ST6, ST149, ST 2538, ST1, ST772, ST97, ST1153, ST672, ST30, ST1535, ST88, ST22, ST8, ST80, ST152, ST1930, and a novel sequence type ST8111 that were assigned in this study table(1). ST5 and ST6 were the predominant clones found in 46% of MRSA isolates (37/80; 46%). Followed by ST88 (9/80; 11.25%), ST22 (7/80; 8.75%), and ST97 (5/80; 6.25%) figure(3). These clones belonged to clonal complexes, CC5 (ST5, ST6, ST149, and ST 2538), CC1(ST1 and ST772), CC97 (ST97 and ST1153), CC361 (ST672), CC30 (ST30), CC15 (ST1535), CC88 (ST88), CC22 (ST22), CC8 (ST8), CC80 (ST80), CC152 (ST152), and CC96 (ST1930). In contrast, there were three MSSA isolates from patients which were assigned to STs: a novel sequence type assigned in this study ST8110, and ST2867 table(1);figure(2.d). The single MSS isolate was *S. argenteus* ST2250.

### SCC*mec* types and *spa* types of *S. aureus* from patients

The most prevalent SCC*mec* types in MRSA isolates were SCC*mec* type IVa, which was observed in (39/80; 48.7%) of isolates. Followed by SCC*mec* type V and its subtype Vc, which was present in (28/80; 35%) of the isolates. Also, SCC*mec* type VI was present in (6/80; 7.5%), as well as SCC*mec* type IV and its subtype IVc were found in (6/80; 7.5%) of the isolates. While SCC*mec* type V/IX was found only in one isolate figure(1). MRSA isolates from patients had 30 *spa* type, t304, t690, t3841, t688, t355, t311, t018, t084, t2297, t127, t4019, t903, t345, t309, t521, t005, t2177, t008, t9736, t045, t267, t4570, t105, t2649, t1627, t4494, t021, t3778, t12438, and t319 (table2). *spa* types t304 and t311 were the most prevalent where t304 accounted for (24/30; 46.6%), and t311 accounted for (13/30; 43.3%) of all the *spa* types among MRSA isolates. Other *spa* types were less prevalent, t688, t690, and t3841 accounting for (6/30; 20%), (5/30; 16.6%), and (5/30; 16.6%), respectively. The remaining *spa* types were distributed among MRSA isolates to a lesser extent. Three MSSA isolates had *spa* types, t159, t2016, and one unknown *spa* type table(2). One MSS isolate, *S. argenteus* had *spa* type t5078.

## Discussion

The emergence and spread of antimicrobial-resistant *S. aureus*, particularly MRSA, is a challenge facing the world today [20]. Between 2000 and 2018, there was an increase in antibiotic consumption in low- and middle-income countries, with the biggest increases in North Africa, the Middle East area, and South Asia [21]. Excessive antibiotic usage in food-producing animals has been linked to drug-resistant bacterial infections in animals and humans [22]. Since the global spread of LA-MRSA began in 2005, many studies have focused on retail meat as a possible transmission environment for MRSA clones [23]. Also, slaughterhouse workers, meat retailers, food handlers, and any individual close to livestock are at risk of acquiring and spread MRSA [24]. The available data on the genetic characterization of *S. aureus*, specifically MRSA in Saudi Arabia, and the genomic relationships of MRSA isolates in human, animal, and retail meat are limited [25, 26]. Here we attempt to close these gaps by performing comparative genomic analysis on Staphylococci and MRSA isolates from humans and meat in terms of occurrence, dissemination, and clonal lineages.

This study identified Staphylococci and MRSA clones isolated from different types of meat (camel, beef, lamb, poultry, and fish) and found 76 Staphylococci of which 11 were MRSA isolates (14.47%), implying that retail food might be contaminated with MRSA and serve as a reservoir of transmission. MLST analysis identified multiple clones found in both meat and patients, indicating the circulation of genotypically diverse *S. aureus* clones. Nearly half of the MRSA isolated from patients and retail belonged to the clonal complex (CC5). Livestock-Associated MRSA (LA-MRSA) clone CC97, was the second dominant clone in meat and had a high prevalence in patients. CC361 is another LA-MRSA clone found in patients and food. Our data demonstrate the presence of CA- and LA-MRSA clones in hospitals. In samples of local and imported retail meat from the Riyadh region, *S. aureus* was found at a prevalence of roughly 24% [26]. Likewise, a previous study revealed that *S. aureus* was present in retail meat at a rate of 25% in Riyadh and had the highest contamination rate with MRSA in camel meat (20%) [25]. Similar to these investigations, this study found that retail meat had a 21.2% *S. aureus* contamination rate, with camel meat having the highest MRSA contamination rate (54%). In a study that analyzed processed food in Riyadh, *S. aureus* prevalence was 62.6% and MRSA prevalence was 56.3% [27]. These rates are to some extent consistent across the country, and considerably higher compared to other Arab countries [28]. The prevalence of MRSA in cattle and cow milk in Algeria, Tunisia, and Jordan was 18%, 20%, and 26% respectively [29–31].

In Saudi Arabia, the MRSA dominant clonal lineage associated with human infections varies from region to region and throughout time. Back in 2015 in Riyadh hospitals, the predominant MRSA clones were CC5, and CC8-ST239-III, and other clones such as CC22-IV, ST30-IV, and CC80-IV. Although the CC8-ST239-III clone was the most prevalent in the GCC region and Saudi hospitals previously, this is no longer the case [32]. Since the emergence of CA-MRSA clones, the pandemic strain CC8-ST239-III has substantially declined but is still sporadically reported in Saudi Arabia [33]. In the eastern region, *Alkharsah* et al. revealed that the most identified clones from infection sites and carriers are CC80 (ST80, ST1440) followed by CC22 (ST22) [15]. Additionally, the most prevalent MRSA lineages among clinical strains in the western region in 2019 were found to be CC5, CC22, CC80, and CC30 [34]. In this study, the most prevalent MRSA lineages in Riyadh city found in raw meat belonged to clonal complexes, CC5 (ST5 and ST6), CC97 (ST97 and ST1153), and CC361 (ST672). While the most prevalent MRSA clones in patients belonged to clonal complexes, CC5 (ST5 and ST6), CC97 (ST97), CC361 (ST672), CC30 (ST30), CC88 (ST88), CC22 (ST22), and CC8 (ST8). Remarkably, none of the clinical MRSA isolates included HA-MRSA clones since all MRSA isolates isolated from patients carried SCC*mec* types IV, IVa, IVc, V, Vc, VI, and IX also the previous predominant clone CC8-ST239-III was not detected.

In this study, the CC5, CC97, and CC361 clonal complexes were shared by MRSA clones from patients and meat. CC5 predominated among MRSA clones in both groups, accounting for (38/80; 47.5%) of the MRSA clones in patients and (5/11; 45.45%) of the MRSA clones in meat. Patients and raw meat shared CC5 (ST5 and ST6) MRSA clones, which carried SCC*mec* types IVa, V, and Vc indicating their belonging to CA-MRSA. Although CC5 is the most prevalent clonal lineage in this study, it was previously a less common clone, making up less than 3% of the isolated clones in a study carried out in Riyadh in 2012 [35]. The second dominant clone from meat in this study was LA-MRSA clone CC97 (ST97 and ST1153), comprising (4/11; 36%) of the MRSA clones from meat, and (6/80; 8%) from patients. Prior to 2018, CC97 clones were also less prevalent in Saudi Arabia, where they made up less than 1.9% in a study conducted in Dammam city and less than 9% of nosocomial infections at a tertiary-care facility in Riyadh [12–15]. Another shared LA-MRSA clone that was detected in both patients and meat is CC361 (ST672). It is represented by (2/11; 18%) of meat MRSA isolates and (5/80; 6%) of patient MRSA isolates. In 2019, this clone first appeared and dominated when it was identified in a tertiary care facility in Riyadh, Saudi Arabia after being reported for the first time in Kuwaiti hospitals one year prior [36, 37].

This study found that the two MSSA clones with the highest prevalence in meat were CC97 (ST1153) accounting for 21.42% and CC361 (ST672) accounting for 16.66% of all the MSSA isolates, both of which are also found in patient MRSA isolates and food. Cattle in Egypt with mastitis were found to have CC1153-MSSA, while in Saudi Arabia, it was identified in humans with skin and soft tissue infections .The *S. aureus* MRSA CC1153 strain is a poorly known Middle Eastern lineage, but it was recently characterized and reviewed in Saudi Arabia, where it was found to harbor several virulent genes, including enterotoxin genes, the penicillinase operon genes, and other antibiotic-resistant genes, implying its ability to cause serious human and livestock infections [38]. Further, MSSA clones, ST361 and ST672 were identified in patients in the western region of Saudi Arabia [34]. According to these findings, meat may be a source of transmission of virulent *S. aureus*, which can result in infections similar to those seen in humans, explaining why common MSSA lineages are prevalent in food and hospitals.

This study identified the two most prevalent MRSA isolates in meat, MRSA ST6-t2450 in camel meat and ST97-t12375 in chicken. However, there is still a lack of information on the *spa* types of MRSA found in meat in Saudi Arabia. In a comparative study conducted in two cities in Saudi Arabia and Egypt in 2015, the most frequent MRSA *spa* type from Saudi Arabia was t008 associated with CC8, which was present also in Egypt but to a lesser extent. Additionally, t084 and t085 *spa* types associated with CC15 were frequent in Saudi Arabia, while in Egypt the most frequent *spa* type was t688 associated with CC5 [39]. This data demonstrates that there has been a shift over time; in this study, MRSA isolates of the t003, t903, and t091 *spa* types predominated in meat, whereas MRSA isolates of the t304, t311, and t688 *spa* types predominated in patients. Furthermore, patients and food MRSA clones shared four *spa* types: t903, t311, t688, and t3841 that suggests that those isolates may have emerged from the same source. Those *spa* types are associated with the following sequence types: ST1153-SCC*mec*IV-t903, ST5-SCC*mec*V-t311, ST5-SCC*mec*V/VI-t688, and ST672-SCC*mec*V-t3841. Similarly, the ST5-SCC*mec*V-t311 and ST5-SCC*mec*VI-t688 genotypes of MRSA were also obtained from goats in Eastern Province, Saudi Arabia, and have been known previously to infect humans [16].

One *S. argenteus* ST2250 isolate was found in each of the patients and meat isolates. This species was previously known as the *S. aureus* clonal complex 75 (CC75) [40]. Locally, a report from 2016 was the first to describe the *S. argenteus* as MRSA lineage CC2250 in a tertiary care facility [12]. Even though it’s now more recognized as a clinically significant species and has recently been reported in many countries, studies have shown that this pathogen has acquired several genes linked with Livestock associated *S. aureus* [40, 41]. *Additionally, six M. sciuri* isolates carrying the *mecA*1 gene were detected in this study in all types of meat except beef. *Mammaliicoccus sciuri*, which is often linked with livestock, carries the *mecA*1 gene, with an overall amino acid sequence similarity of 88% and DNA sequence identity of 80% to MRSA’s *mecA* gene [42]. Out of 30 coagulase-negative Staphylococci (CoNS) isolates, *S*.*sciuri* was detected in 16.66% of meat and dairy products collected in Riyadh, Saudi Arabia [43]. Other CoNS identified in meat in this study, *S. pasteuri, S. saprophyticus*, and *S. haemolyticus*, have received less attention in Saudi Arabia since they are a common component of the human and animal microbiota and an uncommon cause of human illness [44].

The presence of MRSA in raw meat is well-documented worldwide. For instance, in the Czech Republic, (23/65; 35.4%) of the raw meat collected were positive for MRSA, while in the Netherlands the MRSA prevalence in meat was 11.9%, and in the United States, the numbers vary between 3% in some states and 7% in others [4, 45–47]. The occurrence rate of MRSA in meat in this study was 14.47%, which is slightly higher than a previous study that inspected various types of meat in 2016 and reported an MRSA rate of 13% [25]. One limitation of this study is that our sample size was small and planned upon city landscape in a limited geographical area to assess prevalence rates precisely, but our findings are consistent with past research in the region [6, 25, 27]. Another limiting factor of the current study is only MRSA isolates from patients were compared to *S. aureus* isolated from meat products. Future research approaches are needed to look into the presence of virulence and resistance genes in MSSA isolates because even though methicillin resistance is absent, virulence genes may be present and could be a source of pathogenic *S. aureus* for humans. This organism is an important public health concern, and its presence in food, particularly meat, may serve as a potential reservoir for *S. aureus* transmission from livestock to the human population.

## Conclusion

The epidemiology of MRSA and its clonality has shifted as a result of its ongoing genomic evolution. We investigated clonal changes and dissemination of MRSA and MSSA from food and its genetic relatedness to human MRSA isolates. Findings in this study demonstrated that a considerable amount of MRSA and MSSA clones were present in meat products that are shared with those of the patients. Also, these results suggest that the occurrence of CA- and LA-associated MSSA and MRSA, are no longer confined to the community or livestock settings, but have begun to outnumber HA-MRSA in Healthcare settings. These findings might be attributed to a lack of sanitation and veterinary monitoring, as well as the improper use of antimicrobials in the food industry. There is undoubtedly a need for further investigation to identify the distribution, genetic makeup, and antimicrobial resistance of *S. aureus* in food, as well as their effects on human health, furthermore, prescriptive antimicrobial regulations in livestock production, are recommended. It is also suggested that more comprehensive One Health investigations at the local level be performed to better assess the MRSA burden. This study establishes a baseline for Riyadh city, which can help in future developments and comparison for newly implemented policies. A carefully crafted study to track MRSA from livestock, food, and clinic using WGS will be important.

## Ethical Approval

The ethical approval with the number (H-01-R-053) of the study was obtained from The Institutional Review Board (IRB) committee of King Saud Medical City, Riyadh, Saudi Arabia. and 22IBEC051 from Institutional Biosafety and Bioethics Committee of King Abdullah university of science and technology, All procedures were carried out in compliance with the relevant regulations and standards.

## Acknowledgments

This research work was funded by the Saudi Food and Drug Authority (SFDA), Riyadh, Saudi Arabia, and King Abdullah University of Science and Technology (KAUST), Thwal, Saudi Arabia. The authors would like to thank the University of Jeddah for their support.

## Declaration statement

The views expressed in this paper are those of the authors and not do not necessarily reflect those of the SFDA or its stakeholders. Guaranteeing the accuracy and the validity of the data is a sole responsibility of the research team.

## Supporting information

### data availability Data availability

Sequence data for this work are deposited in the ENA under accession number PRJEB64197.

